# Pheno-RNA, a method to identify genes linked to a specific phenotype: application to cellular transformation

**DOI:** 10.1101/2020.03.04.976910

**Authors:** Rabih Darwiche, Kevin Struhl

## Abstract

Cellular transformation is associated with dramatic changes in gene expression, but it is difficult to determine which regulated genes are oncogenically relevant. Here, we describe Pheno-RNA, a general approach to identify candidate genes associated with a specific phenotype. Specifically, we generate a “phenotypic series” by treating a non-transformed breast cell line with a wide variety of molecules that induce cellular transformation to various extents. By performing transcriptional profiling across this phenotypic series, the expression profile of every gene can be correlated with the transformed phenotype. We identify ~200 genes whose expression profiles are very highly correlated with the transformation phenotype, strongly suggesting their importance in transformation. Within biological categories linked to cancer, some genes show high correlations with the transformed phenotype, but others do not. Many genes whose expression profiles are highly correlated with transformation have never been linked to cancer, suggesting the involvement of heretofore unknown genes in cancer.

---

A variety of whole-genome approaches have been employed to determine which genes are important for a specific phenotype. Transcriptional profiling of cells under the relevant developmental or environmental conditions identifies differentially expressed genes^1–4^. While some of these are likely important for the process of interest, it is unlikely that all of them are. In this regard, there is surprisingly limited conservation of genes in different yeast species that respond to a given stress^5^. Alternatively, classical or systematic genome-scale mutational screens identify genes that affect a given phenotype^6–8^. However, mutations might confer such phenotypes by indirect and/or artifactual effects, and many important genes might be missed due to the nature of the assay or to functional redundancy. Integration of other forms of information (e.g. genome binding, chromatin status, protein-protein interactions, genetic epistasis, evolutionary conservation) is very helpful in addressing the connection between genes and phenotypes^9^. Here, we describe a new genome-scale approach, Pheno-RNA, to identify genes important for a specific phenotype.

We apply Pheno-RNA to the process of cellular transformation, using our inducible model in which transient activation of v-Src oncoprotein converts a non-transformed breast epithelial cell line into a stably transformed state within 24 hours^10, 11^. This epigenetic switch between stable non-transformed and transformed states is mediated by an inflammatory positive feedback loop involving NF-κB, STAT3, and AP-1 factors^11, 12^, and many genes are directly and jointly co-regulated by these factors^13^. In addition, we identified > 40 transcription factors important for transformation as well as putative target sites directly bound by these factors^14^. These transcriptional regulatory circuits are utilized in other cancer cell types and human cancers, and they are the basis of an inflammation index to type human cancers by functional criteria^14^. Nevertheless, and despite the molecular description of these regulatory circuits, it is unclear which genes are important for transformation.

Pheno-RNA is designed to distinguish genes that are “drivers” as opposed to “passengers” for a specific phenotype. Cells are subject to a wide variety of experimental perturbations, and the expression profiles of individual genes are correlated with a quantitative measurement of a given phenotype. The specific perturbations, individually or in combination, do not matter for Pheno-RNA analysis, although they can provide additional information. In principle, the expression profiles of genes driving the phenotype should be highly correlated with the degree/strength of the phenotype. Conversely, genes whose expression is regulated by one or a few experimental conditions are likely passengers under those conditions and not generally relevant for the phenotype.

To identify genes important for breast cellular transformation, we generated a series of transformation phenotypes by treating the parental MCF-10A cell line (i.e. lacking the ER-Src derivative) with a wide variety of signaling molecules - interleukins, growth factors (RGF, FGFs, HGF, TGFs), TNF-α, IFN-γ, leptin, insulin, TPA, cAMP - either individually or in combination. The degree of transformation was determined quantitively by growth in low attachment, an assay highly correlated with the more qualitative soft-agar assay^15^. With the exception of Oncostatin M, none of signaling molecules tested individually caused transformation. Out of hundreds of combinations tested (**Supplementary Table 1**), we selected 17 to generate the phenotypic series. These 17 treatments show a wide range of transformation phenotypes (**Fig. 1a**), but do not significantly affect cell growth in standard (high attachment) conditions (**Fig. 1b**).

**Fig. 1.**
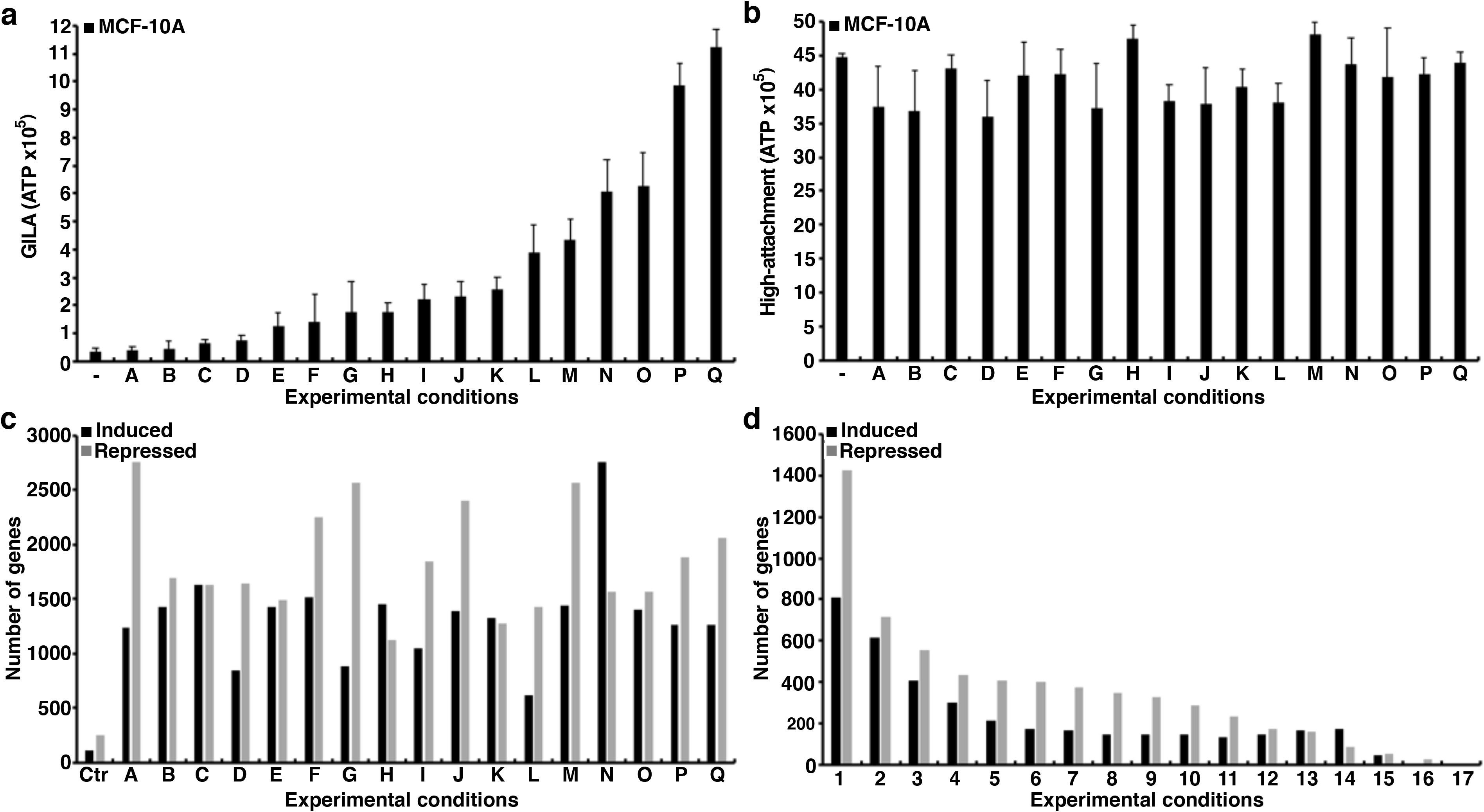
Transformation levels and differentially expressed genes. **(a, b)** Relative growth levels of MCF-10A cells grown on low-attachment (**a**) or high-attachment (**b**) plates for 5 days in 17 different conditions (A-Q; see **Supplementary Table 1)**. **(c)** Number of genes whose expression is induced (black) or reduced (gray) 2-fold under the 17 conditions. The control represents the absence of any compound added to the cells. **(d)** Number of genes whose expression is induced or reduced under the indicated number (1-17) of experimental conditions.

We performed RNA-seq experiments to obtain transcriptional profiles under these 17 conditions (**Supplementary Table 2**). As expected, a few thousand genes were up- or down-regulated (> 2-fold) in each condition as compared to the untreated control (**Fig. 1c**), and some genes were up- or down-regulated under multiple conditions (range from 1-17; **Fig. 1d**). For 6651 genes, we determined the Pearson’s correlation coefficient between the mRNA levels and degree of transformation under the 17 experimental conditions (**Supplementary Table 3**). The correlation coefficients range from 0.99 to −0.60, as compared to the ± 0.4 range observed when mRNA values are randomized among the different conditions (**Fig. 2a**). 209 and 82 genes have remarkably high positive correlation coefficients (> 0.8 and > 0.9, respectively), strongly suggesting their relevance for cellular transformation. The results for negatively regulated genes are less dramatic, but 52 genes have correlation coefficients < −0.5. Importantly, for any individual condition, only a subset of genes (range 0.5 to 5.5%) showing regulated expression have high correlations (> 0.6) with the transformation phenotype.

**Fig. 2.**
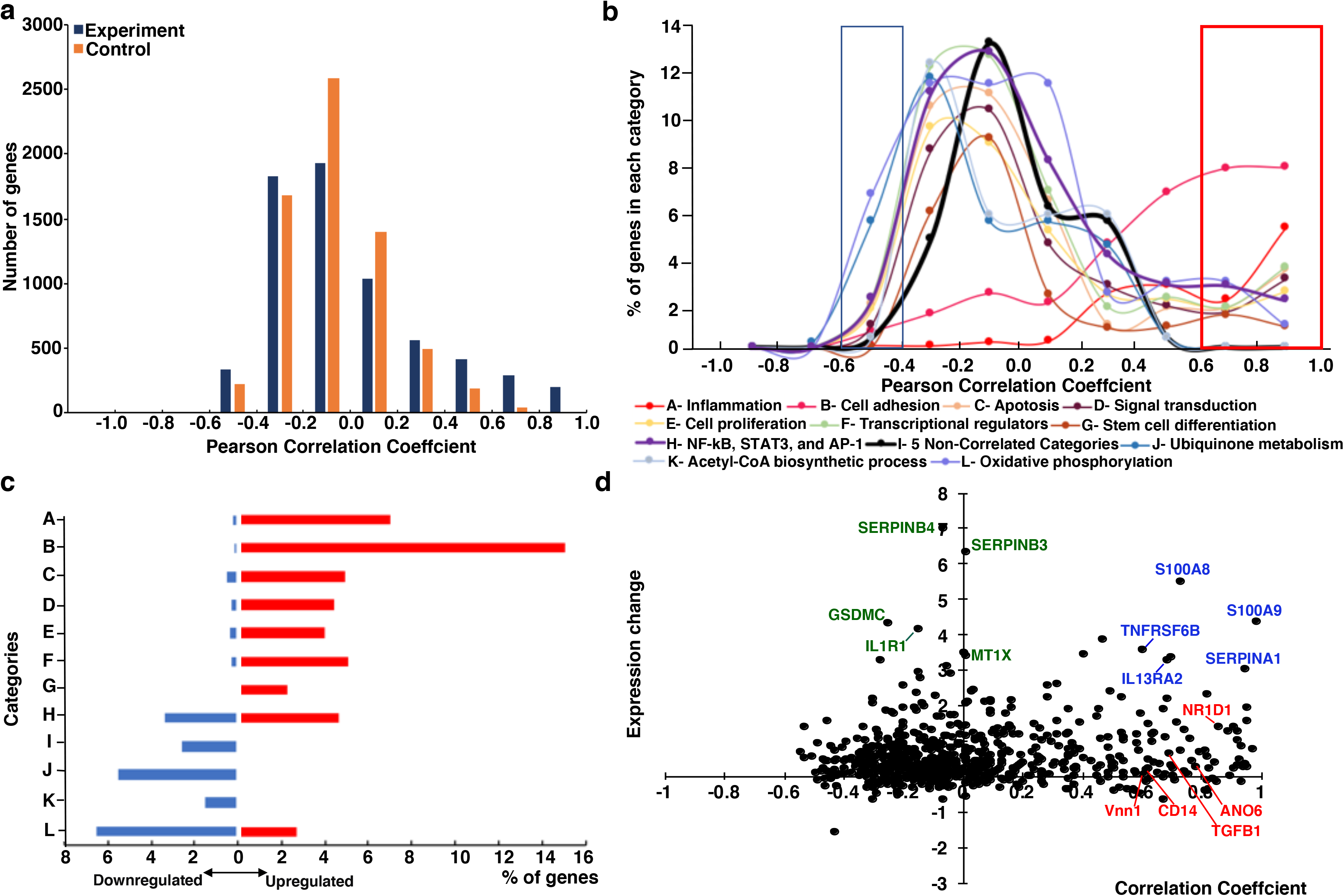
Correlation of gene expression profiles to transformation phenotype. **(a)** Number of genes in the indicated bins of correlation coefficients in the experimental (purple) and shuffled (orange) RNA samples. (**b**) Percent of genes in each functional category (A-L shown in various colors) in the indicated binds of correlation coefficients. The red box indicates high positive correlation (> 0.6) and the blue box indicates high negative correlation (< 0.4). **(c)** Percent of genes in the indicated category (A-L as above) that have high positive (> 0.6; red) or negative (< 0.5; blue) correlation to the level of transformation. **(d)** Relationship between the correlation coefficient and fold-induction upon Src induction (log 2) for all genes (black dots) that are part of the NF-κB/STAT3/AP-1 transcriptional network. Selected inflammatory genes with high correlation/high induction (blue), low correlation/high induction (green), and high correlation/low induction (red) are indicated.

**Fig. 3.**
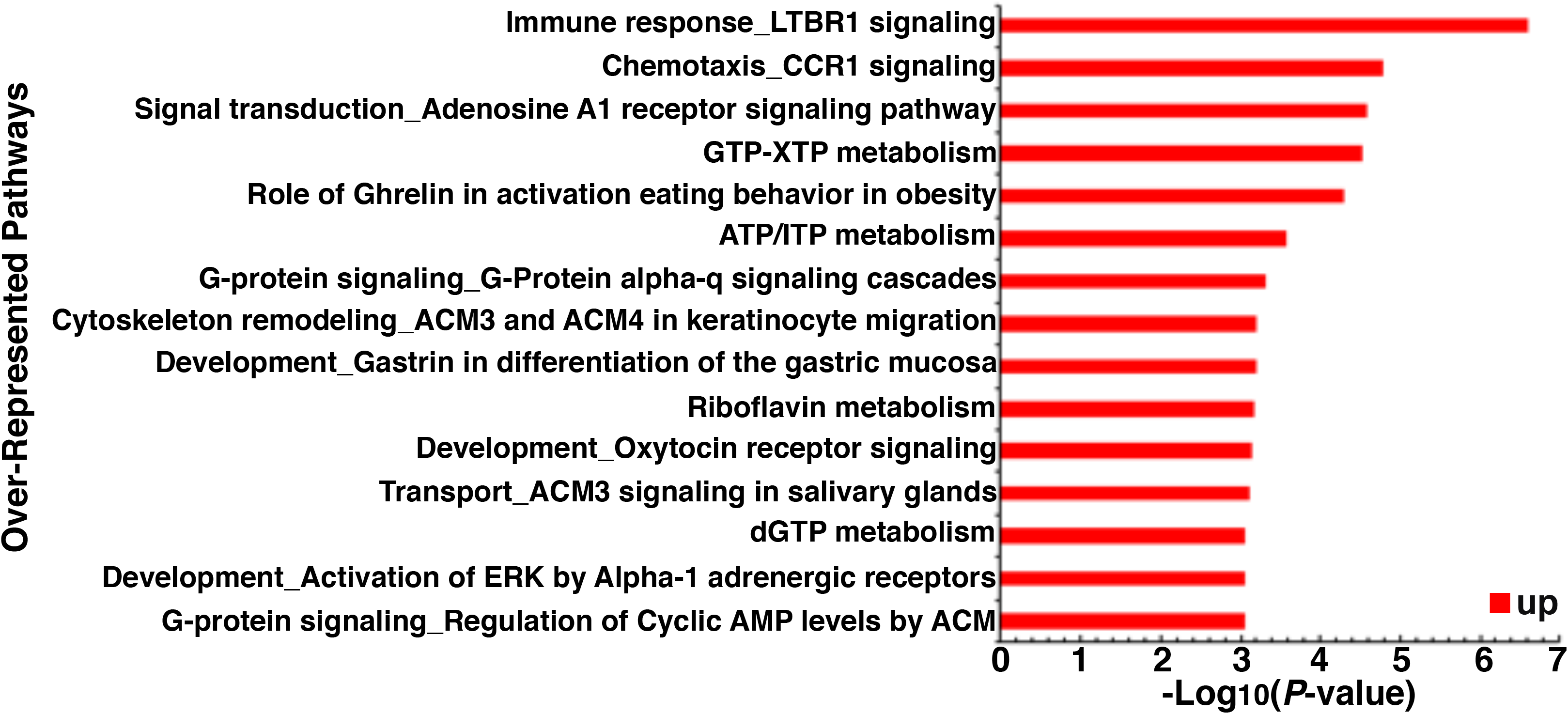
Over-represented gene categories of 90 genes with high correlation (> 0.8) to the transformed phenotype but no previous link to cancer. The significance of each category is indicated by −Log_10_(*P*-value).

As expected, genes whose expression profiles are highly correlated (either positively or negatively) with transformation are enriched in pathways strongly associated with cancer progress such as the inflammatory response, apoptosis, metastasis, and transcriptional regulators (**Fig. 2b, c; Supplementary Table 4**). In addition, genes showing strong correlation (> 0.8) with the transformation phenotype are enriched for genes that are up- or down-regulated in the Src-inducible model and/or are part of the transcriptional regulatory network mediated by the combined action of NF-κB, STAT3, and AP-1 (**Fig. 2b, c**)^13, 14^. However, 76% of these high-correlated genes are not regulated in the Src-inducible model of transformation 85% are not part of the transcriptional regulatory network. Thus, Pheno-RNA both confirms genes and pathways previously linked to the transcriptional phenotype and also identifies genes that were not identified by other approaches.

Although the inflammatory pathway is critical for transformation, our results distinguish among inflammatory genes whose expression patterns are strongly correlated with the transformation phenotype from those that are not (**Fig. 2d**). Some inflammatory genes that are induced by Src have expression levels that are highly correlated with the transformation phenotype (examples in blue), whereas other Src-inducible genes show essentially no correlation with the transformation phenotype (examples in green). Conversely, other inflammatory genes have expression profiles highly correlated with transformation, but are not induced in our ER-Src model (examples in red). These results suggest a subset of inflammatory genes that are important for transformation. Similar analyses on non-inflammatory pathways previously linked to transformation (e.g. apoptosis, cell proliferation, and metastasis) also reveal subsets of genes whose expression patterns strongly correlate with transformation.

As Pheno-RNA is based solely on the connection between specific phenotypes and gene expression patterns in a defined experimental system, it represents a new and unbiased approach to identify genes associated with transformation. We therefore used the following approach to identify genes whose expression patterns are strongly correlated to the transformation phenotype, yet have never previously linked to cancer. First, we performed an automated PubMed search (NCBI’s E-Utilities) on each gene ID and asked whether it was it was linked to studies on PubMed. Second, we reviewed the literature to confirm that the candidate cancer genes were not linked to cancer. Third, we cross-checked the candidate genes against the Network of Cancer Genes (NCG) that contains detailed information on 2,372 cancer genes, the Cancer Gene Census (CGC) database that catalogues genes with mutations causally implicated in cancer, Tumor-Associated Genes (TAG), and COSMIC and cBIOPortal that focus on cancer alterations rather than cancer genes.

From this approach, we identified 90 genes that are strongly linked to the transformation phenotype (R > 0.8) but not previously linked to cancer (**Supplementary Table 3**). Interestingly, these genes are enriched in functional categories including the ribosome, translational elongation, and RNA transport. Some of these 90 genes are up-regulated in the Src-inducible model, whereas others are not. Conversely, we identified 6 genes negatively linked to the transformation phenotype (R < −0.5) and previously unlinked to cancer. Lastly, expression of many small nucleolar RNAs (A45B, D36B, D18A, D7A, D42B, D90, and D49B) show strong correlations (> 0.6) with the transformed phenotype, yet have never been linked to cancer. These genes are new candidates for playing a role in cancer, and hence worthy of functional analysis, either individually or in combination.

Our Pheno-RNA analysis distinguish between genes that are strongly associated with cellular transformation from those that are regulated by a specific condition that causes a transformation phenotype, and it also identifies new genes that are candidates for affecting the phenotype. Presumably, genes with high correlation coefficients are important for the transformation phenotype assayed here (growth under conditions of low attachment), whereas regulated genes with poor correlation coefficients are more relevant for a non-oncogenic function(s) of the specific experimental condition(s). However, it remains possible that some of these low-correlating genes are important for the transformation phenotype, particularly if pairwise (of higher order) interactions between genes affect their transcriptional profiles and oncogenic function.

More generally, Pheno-RNA can be applied to multiple phenotypes generated by the experimental conditions. For example, the experimental conditions and transcriptional profiling data presented here can be linked to other transformation-related phenotypes such as morphology, focus formation, cell motility/invasive growth, mammosphere formation, and metformin sensitivity. The phenotypic series will likely differ among these various phenotypes, making it possible to identify different sets of genes associated with these phenotypes. Lastly, Pheno-RNA should be generally applicable for any phenotype that can be quantitated and for which multiple experimental conditions can be used to generate a phenotypic series.

## MATERIALS AND METHODS

The non-transformed breast cell line MCF-10A^16^ was cultured as described previously^10, 11^. The level of transformation mediated by compounds listed in Table were determined by growth in low attachment (GILA assay) essentially as described previously^15^ except that 384-well plates were used and the cell concentration was optimized to 25 cells per well in 30 μl. Cells were assayed for ATP content after 5 days of incubation at 37°C, with each plate having five repetitions, and the entire screen repeated three times.

Transcriptome level mRNA profiling was performed by 3’READS as described previously^17^. For each condition, reads counts for each gene were normalized to the total number of True Poly(A) Read pairs. Normalized mRNA seq values for the 17 different experimental conditions (Supplementary Table 2) were correlated to the level of transformation using the Excel CORREL function. As a control, mRNA seq values for every gene were randomly shuffled across the 17 treatments and then correlated to the transformation level. Over-representational significance (*p*-value) for each set of genes was calculated using the Fisher Exact test.

Gene sets based on correlation cutoffs were used as input for MetaCore^TM^ Functional Enrichment by Ontology using pathway maps in order to identify biological pathways associated with transformation (https://portal.genego.com/cgi/data_manager.cgi). An FDR cutoff of 0.05 was used to identify pathway maps significantly associated transformation. Network analysis were also performed using Gene Set Enrichment Analysis (GSEA) and DAVID ontology.

## ACKNOWLEDGEMENTS

We thank Zarmik Moqataderi and Joseph Geisberg for help with the bioinformatic analysis and useful discussions. This work was funded by the Swiss National Science Foundation SNSF (Project P2FRP3_178090 to RD) and by a research grant to K.S. from the National Institutes of Health (CA 107486).

